# Transcutaneous Vagus Nerve Stimulation Reduces Pain and OA Progression in Mouse Models of Post-Traumatic Osteoarthritis

**DOI:** 10.1101/2024.10.31.621385

**Authors:** Shivmurat Yadav, Lynsie Morris, Taylor Conner, Jessica Lumry, Sanique M South, Emmaline Prinz, Vladislav Izda, Gabby Dyson, Montana Barrett, Stavros Stavrakis, Matlock A Jeffries, Timothy M Griffin, Mary Beth Humphrey

## Abstract

Currently, there are no disease-modifying osteoarthritis (OA) drugs (DMOAD) to prevent OA progression and there are limitations on pain relieving therapeutics. Vagus nerve stimulation (VNS) delivered by an implantable device is FDA-approved for refractory epilepsy and severe depression. Here, we investigated the efficacy of transcutaneous VNS (tVNS) for preventing OA progression and providing pain relief in two mouse models of post-traumatic OA (PTOA): the surgical destabilized medial meniscus (DMM) and the non-surgical forced tibial compression anterior cruciate ligament rupture (ACLR). Here, we show that 2 weeks of tVNS significantly reduced histological OA scores in male and female mice after ACLR compared to sham stimulation. In female, but not male, mice, tVNS reduced hyperalgesia and mechanical allodynia. In the slower DMM model, 8 weeks of tVNS improved weight bearing in male and female mice, but only female mice had improved hyperalgesia. Male mice had lower OA histological scores. Serum proinflammatory cytokines were significantly reduced by tVNS in both models but differed by gender and model. Overall, these results provide strong pre-clinical evidence that tVNS reduces OA progression, improves pain, and suppresses pro-inflammatory cytokines, making it a promising DMOAD.

## Introduction

Historically, OA was not considered an inflammatory disease, but recent studies show low-grade synovitis is common and occurs in a substantial portion of patients (41% vs 14% without OA)(1). Low-grade inflammation correlates with OA severity, including immune cells in the synovium, macrophage infiltration of the infrapatellar fat depots, and pro-inflammatory cytokines(2, 3). In human OA, *in vivo* analysis of activated macrophages using etarfolatide labeling and SPECT-CT reveals that 76% of OA knees have an accumulation of activated macrophages that are significantly associated with knee pain (R=0.60, p<0.0001), joint space narrowing (R=0.68, p=0.007), and osteophytes (R=0.66, p=0.001)(1). These activated macrophages are thought to respond to danger-associated molecular patterns (DAMPs) and secrete pro-inflammatory cytokines like TNFα, IL-1β, and IL-6. Additionally, these macrophages interact with T cells, the second most abundant cell type in synovitis(4). Inflammation likely triggers a cascade of inflammatory mediators, including proteases, prostaglandins, cytokines, lipids, chemokines, and neuropeptides, contributing to pain and joint destruction(5).

Currently, there are no disease-modifying drugs to prevent OA progression and only limited options to treat OA-associated pain. A poor understanding of key features predicting OA progression and pain mechanisms involving nociceptive (inflammation and damage), neuropathic (nerve damage), and centralized (disturbance in central nervous system pain and processing) have prevented the development of effective therapies to limit pain and OA progression. Vagus nerve stimulation (VNS) has emerged as a powerful method to induce systemic anti-inflammatory effects and improve outcomes in epilepsy(6), heart failure(7), and rheumatoid arthritis in humans and rodent models(8, 9). VNS stimulation of the parasympathetic nervous system is postulated to have three modes of action. Firstly, VNS stimulates the hypothalamic-pituitary-adrenal axis and induces the release of glucocorticoids(10). Secondly, VNS stimulates splenic T cells to produce choline acetyltransferase (ChAT), releasing Ach and stimulating splenic macrophages through the nicotinic acetylcholine receptor α7 (α7nAchR) to inhibit the production of pro-inflammatory cytokines, thus reducing systemic inflammation(11, 12). Thirdly, within the joint, cholinergic fibers innervate the synovium, trabecular bone, and periosteum, and studies suggest that the parasympathetic nervous system modulates nociceptive pain and possibly OA pathogenesis(13). Transcutaneous VNS (tVNS) exerts anti-inflammatory effects similar to cervical VNS in rodent models and humans(7, 14).

In this study, we hypothesized that tVNS would decrease pain and limit the histological progression of OA by reducing inflammation or modulating the activities of neurons in the pain pathways. We employed two preclinical mouse models of knee OA and applied tVNS or sham stimulation to mice after knee joint injury. Our results show that tVNS reduced a subset of pain measures in both models, with more significant effects seen in female mice. tVNS improved lateral joint compartment histological OA scores in DMM and ACLR models and the medial compartment in the ACLR model. tVNS reduced several pro-inflammatory cytokines in a sex and model-dependent manner. This work is significant as it provides crucial preclinical evidence supporting the efficacy of non-invasive tVNS to treat OA pain and reduce OA progression by suppressing inflammation, thus providing a strong rationale for randomized clinical trials of tVNS in human OA.

## Methods

### Ethics statement

All experimental procedures were approved by the Institutional Animal Care and Use Committees at Oklahoma Medical Research Foundation (OMRF) and Oklahoma City VA, and experiments were performed in accordance with the ethical guidelines.

### Animals and Experimental Design

C57BL/6J male and female mice were purchased from the Jackon Laboratory (Bar Harbor, ME USA), acclimatized, and underwent forced tibial compression to induce ACL rupture of the right leg (15–17) or DMM surgery at 16 weeks old. After ACL-rupture, mice were randomized to receive 2 weeks of either tVNS or SHAM stimulation for 10 minutes a day, 5 days a week (Fig. 1A). Electrical stimulation was administered using a transcutaneous electrical nerve stimulation device (InTENSity Twin Stim; Current Solutions LLC, Austin, TX), for 10 minutes per day, 5 days per week, under 2% isoflurane anesthesia. In the active tVNS group, electrodes were positioned over the auricular concha region with the cathode inside and the anode outside, delivering a 2-mA current, as previously described (7). In the sham group, electrodes were placed similarly, but the device remained off. Behavioral and pain measurements were taken one day before the ACL injury (baseline) and two weeks after tVNS or sham treatment (Fig. 1A). After DMM surgery, mice were housed for 4 weeks to develop early OA before randomization to receive 8 weeks of tVNS or sham stimulation (Fig.1H). Behavioral and pain measurements were performed before treatment and after 4 and 8 weeks of tVNS or sham stimulation (Fig. 1H). Animals were euthanized after 2 weeks (ACL rupture) or after 12 weeks (DMM), and knee joints and blood were collected (Fig.1A&H).

**Figure 1.**
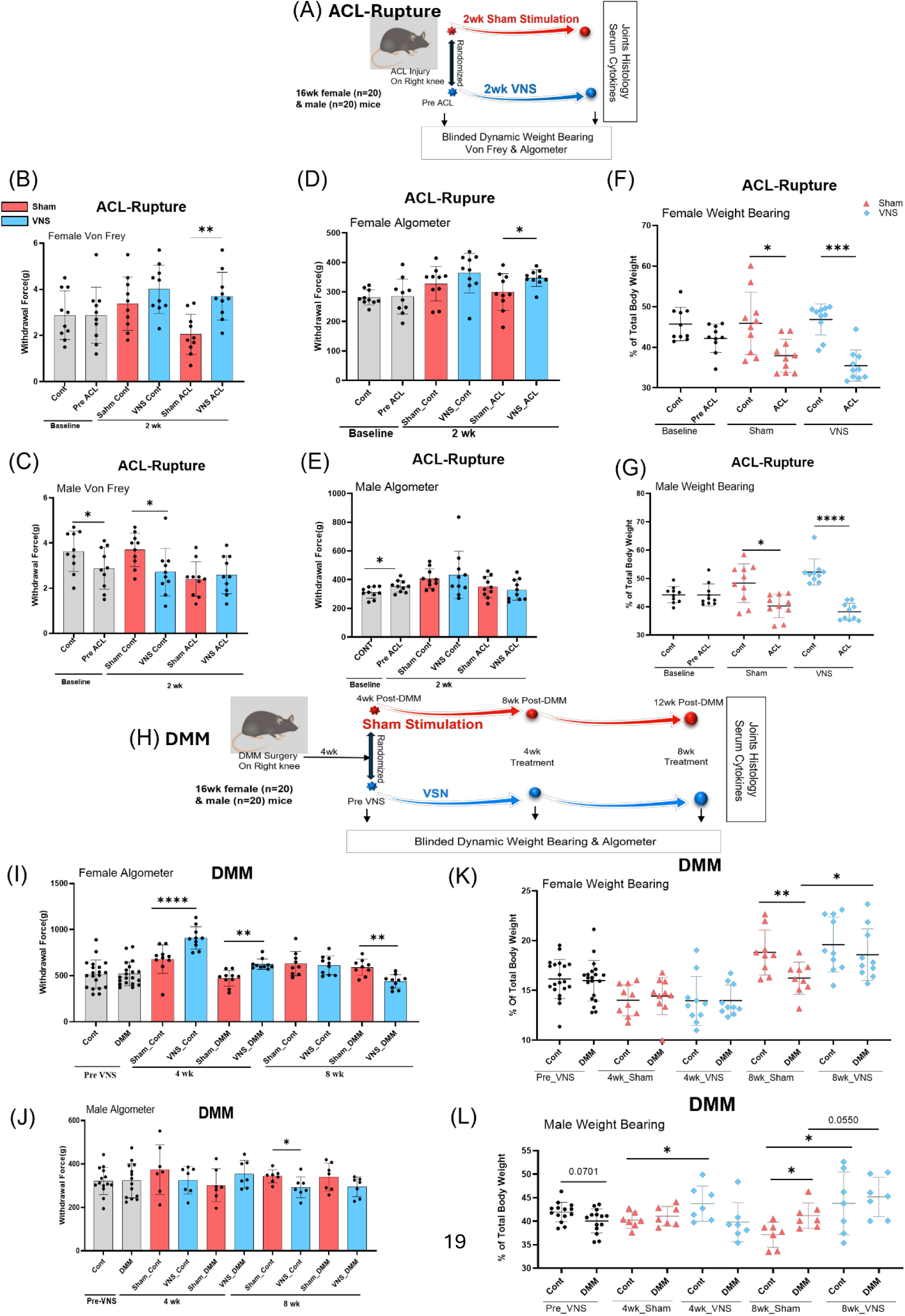
tVNS reduces acute and chronic knee pain and improves weight bearing. (A) Forced tibial compression ACL-rupture study design. (B & C) von Frey withdrawal force in ACL females and males at baseline and after 2 weeks of treatment. (D & E) Algometer withdrawal force of the contralateral and ACL-rupture knees of females and males. (F & G) Percentage of weight bearing on the contralateral and ACL-rupture limbs in females and males. (H) DMM study design. (I & J) Algometer withdrawal force on the contralateral and DMM knees before tVNS (PreVNS) and after 4 or 8 weeks of treatment in females and males. (K & L) Percentage of weight bearing on the contralateral and DMM limbs in female and male mice. Data are presented as mean ± SD, with n = 7 to 10; *p < 0.05, **p < 0.01, ***p < 0.001, ****p < 0.0001. Statistical analyses involved two-tailed paired t-tests or unpaired t-tests to compare two groups as applicable. Abbreviations: Cont = Contralateral, DMM = Destabilized Medial Meniscus, ACL = Anterior Cruciate Ligament, tVNS = transcutaneous Vagus Nerve Stimulation, SHAM = SHAM tVNS. Red is SHAM treated; Blue is tVNS treated.

### Spontaneous Pain Evaluation: Dynamic Weight-Bearing

Spontaneous pain in the hind limbs (rear left and rear right leg) was evaluated using a Dynamic Weight-Bearing (DWB) System apparatus (BIOSEB) according to BIOSEB’s instructions. After a 2-minute latency period, the software automatically recorded two 5-minute acquisition sessions for each mouse. The two readings were then averaged to represent the weight exerted by each hind limb. The weight distribution between the right and left paw was calculated as a percentage of the total body weight.

### Mechanical Hyperalgesia: Algometer

Mechanical hyperalgesia was measured using the Bioseb SMALGO (Small Animal ALGOmeter) according to the manufacturer’s instructions. A higher force threshold indicates lower perceived pain levels in the animal, whereas a lower force threshold indicates higher pain sensitivity. To ensure reliable measurements of mechanical sensitivity, the test was performed three times on each knee, with a 10-minute waiting period between tests; the three values collected for each knee were then averaged and analyzed to compare the sensitivity thresholds between the groups. The threshold was expressed as the force (in grams) that elicited a response. The evaluation was performed in a blinded fashion.

### Mechanical Allodynia Evaluation: Digital Von Frey

The electronic Bioseb Von Frey filament was utilized to determine the mechanical sensitivity threshold in mice according to the manufacturer’s instructions. Mice were habituated for 45 minutes in the Von-Frey stand. The withdrawal threshold was determined three times on each hind paw. Values were expressed as the force (in grams), where a higher force threshold indicates lower perceived pain levels in the animal, while a lower force threshold indicates higher pain sensitivity. The evaluation was performed in a blinded fashion.

### Mice euthanasia and sample collection

Blood samples were collected by cardiac puncture at the time of euthanasia. Serum samples were stored at −80 °C. Injured and contralateral knees were collected, and muscle removed.

### Knee Histology

Knees were fixed in 4% formalin and decalcified in 10% ethylenediaminetetraacetic acid (EDTA) (pH 7.2-7.4) for 14 days at 4°C. Knees were paraffin-embedded, followed by sagittal sectioning for ACL rupture knee and coronal sectioning for DMM surgery knee. Sections were stained with Safranin-O/fast green stain and counterstained with hematoxylin. The DMM histology was evaluated using the OARSI grading method by two blinded graders, while ACL rupture histology was evaluated using both the OARSI grading system and an anterior versus posterior zone-based semi-quantitative OA scoring system by three blinded graders(18).

### Serum cytokines analysis

Serum cytokines, including IL-1α, IL-1β, IL-2, IL-3, IL-4, IL-5, IL-6, IL-9, IL-10, IL-12 (p40), IL-12 (p70), IL-13, IL-17A, Eotaxin, G-CSF, GM-CSF, IFN-γ, KC, MCP-1 (MCAF), MIP-1α, MIP-1β, RANTES, and TNF-α, were measured using the Bio-Plex Pro Mouse Cytokine 23-plex Assay kit (BIO-RAD, #M60009RDPD). Each mouse sample was tested in duplicate.

### Statistical analysis

Statistical analysis was performed using GraphPad Prism software (version 10.2.6). Animal group sizes were determined by power analysis using algometer data as the primary outcome from our pilot study, estimating that 9 animals per group would provide 80% power to detect an effect size of 1.26 at a significance level of p = 0.05. For comparing two groups, either paired or unpaired t-tests or Mann-Whitney test with two-tailed analysis and 95% confidence intervals were used to assess mean differences, as applicable. A two-way ANOVA followed by Fisher’s LSD multiple comparison test was employed to evaluate the effects of two factors. Simple linear regression analyses explored relationships between variables using correlation coefficients and regression slopes. Detailed descriptions of statistical significance (p-values) and animal numbers per group are provided in figure legends.

## Results

### tVNS treatment improves mechanical allodynia and hyperalgesia in female mice after ACL rupture

To determine the effects of tVNS treatment on acute post-traumatic knee pain, we used the forced ACLR model (Fig. 1A). Age and sex-matched mice underwent ACL rupture of the right leg. Both sexes were included to determine whether tVNS stimulation would have sex-dependent effects. The contralateral leg served as the control uninjured knee joint. The day after rupture, mice were randomized to tVNS- or sham-tVNS treatment for 10 minutes daily, five days a week, for two weeks. Three pain outcomes were measured before the injury and after two weeks of tVNS or sham stimulation: digital von Frey assessed mechanical allodynia, pressure algometry measured mechanical hyperalgesia, and weight-bearing analysis to determine the weight on the injured limb. The study was concluded on post-rupture day 15.

Compared to the contralateral leg, von Frey analysis showed significantly reduced withdrawal force in sham-stimulated ACL-injured limbs, indicating that ACLR model induced mechanical allodynia (Fig. 1B and 1C, red bars). tVNS treatment significantly improved the withdrawal force in female mice but not male mice, indicating that tVNS improved mechanical allodynia in female mice (Fig. 1B and 1C, blue bars). Pressure algometry analysis showed decreased withdrawal force in ACL-injured sham-stimulated female mice, and it was significantly increased by tVNS treatment (Fig. 1D). Male mice had no differences in withdrawal force between the ACL-ruptured and contralateral limbs at the 2-week time point independent of tVNS or sham stimulation (Fig. 1D and 1E). These results indicate that tVNS treatment improves mechanical allodynia and hyperalgesia in a sex-dependent manner.

### tVNS treatment improves mechanical hyperalgesia in female mice with DMM-induced PTOA

Next, we used the DMM surgery model of PTOA to test the efficacy of tVNS on chronic OA pain (Fig. 1H). Age and sex-matched adult mice underwent DMM surgery on the right leg with no intervention for 4 weeks post-surgery, allowing early OA to develop. Before the randomization of mice on postoperative day 30, pressure algometry and weight-bearing analysis were performed. After cage randomization, mice received tVNS or sham stimulation 5 days a week for 8 weeks as indicated above. Pain measurements were repeated 4 and 8 weeks into treatment. The study was concluded at postoperative week 12.

Algometer withdrawal thresholds were similar between the DMM and contralateral leg at postoperative week 4 for both sexes before randomization (Fig. 1I and 1J). After 4 weeks of stimulation, female mice receiving tVNS treatment had improved withdrawal thresholds compared to sham, indicating that tVNS treatment improved mechanical hyperalgesia (Fig. 1I). Male mice had a trend towards improvement in hyperalgesia (Fig. 1J) After an additional 4 weeks of tVNS treatment, female mice developed significantly lower withdrawal responses than sham-treated mice, suggesting that prolonged tVNS treatment might potentiate mechanical hyperalgesia. In male mice, tVNS treatment failed to improve mechanical hyperalgesia. (Fig. 1J) These results indicate that short-term therapy with tVNS improves hyperalgesia in chronic knee OA, but the effect is more significant in female mice.

### tVNS treatment improves weight-bearing in DMM-but not ACLR-induced PTOA

To assess the effects of tVNS on acute or chronic knee pain-associated behavior, we analyzed dynamic weight bearing in our mouse models of early and late knee OA. Two weeks after ACL injury, female and male mice developed significant weight-bearing abnormalities on their injured limbs, with significantly more weight-bearing on their contralateral limb (Fig. 1F and 1G). tVNS treatment failed to improve weight bearing compared to sham-treated mice. In the DMM model, female mice developed significantly decreased weight bearing on the DMM limb at postoperative week 12 (Fig. 1K). Compared to sham treatment, 8 weeks of tVNS improved weight bearing on the DMM limb (Fig. 1K). Male mice receiving 8 weeks of tVNS treatment also had improved weight bearing on the DMM limb compared to sham stimulation (Fig. 1L). These data indicate that 8 weeks of tVNS treatment improves pain associated with bearing weight in the chronic DMM-induced OA model in both sexes but not in the ACLR model.

### tVNS effects on joint damage in PTOA

Two weeks after ACLR, male and female mice demonstrated significant OA histological changes in the operated knee (Fig. 2A), with elevated OARSI scores and anterior and posterior zone-based semi-quantitative OA scores (Fig. 2B-E). Compared to sham stimulation, tVNS treatment significantly reduced OARSI and anterior zone OA scores of the medial femur compartment of male and female mice. (Fig. 2B & 2D). There was no improvement in OA scores with tVNS in the other medial tibia or lateral knee compartments. (Fig. 2B-E).

**Figure 2.**
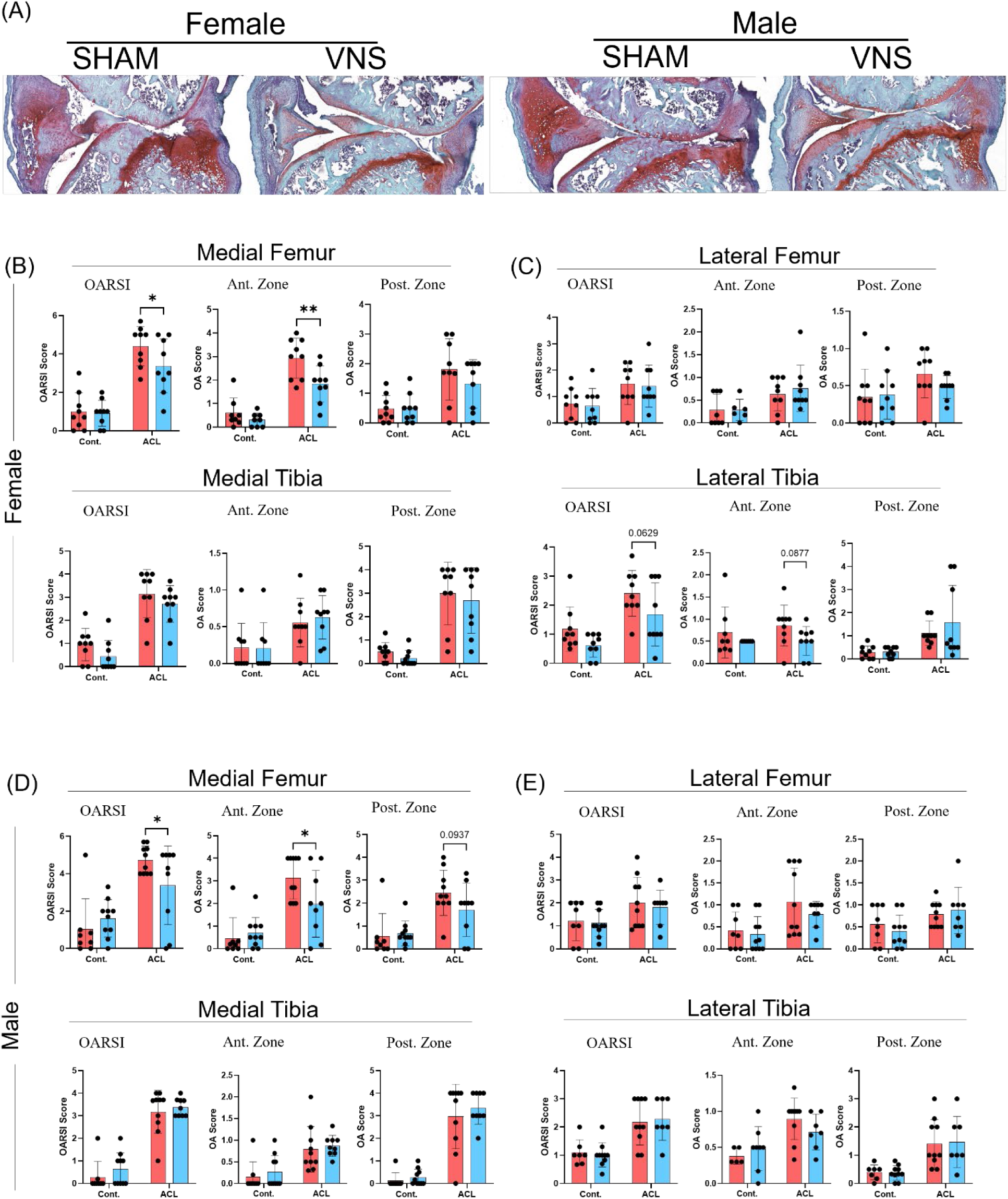
tVNS reduces PTOA after ACL rupture. (A) The top panel displays representative sagittal histology sections of ACLR joints. The bottom panels show OARSI and OA scores by compartment, with sham stimulation in red and tVNS in blue. Scores are included for non-injured contralateral knees and ACL-ruptured knees. (B) Female medial femur or tibia; (C) Female lateral femur or tibia; (D) Male medial femur or tibia; (E) Male lateral femur or tibia. Data are presented as mean ± SD, with n =9 to 10; *p < 0.05, **p < 0.01. Two-way ANOVA followed by Fisher’s LSD multiple comparisons test was used to evaluate the effects of injury and treatment in the statistical analyses. Abbreviations: Cont = Contralateral, DMM = Destabilized Medial Meniscus, ACL = Anterior Cruciate Ligament, VNS = Vagus Nerve Stimulation, SHAM = SHAM VNS Treatment, Ant. Zone = Anterior Zone, and Post. Zone = Posterior Zone. Red is SHAM treated; Blue is tVNS treated.

At post-DMM week 12, male mice developed mild OA as indicated by OARSI scores compared to females (Fig. 3 and S1). Safranin-O/fast green stained images of articular cartilage and subchondral bone showed damaged medial meniscus in the DMM knee, confirming successful DMM surgery in both males and females (Fig. 3A and S1A). In male mice, 8 weeks of tVNS treatment showed significant improvement in the OARSI scores of the lateral femur compartment compared to sham treatment (Fig. 3C). No significant improvements were observed in the medial compartments of the tibia and femur (Fig. 3D and E). In females, OARSI scores were low and tVNS did not modify the mild OA findings (S1B-E).

**Figure 3.**
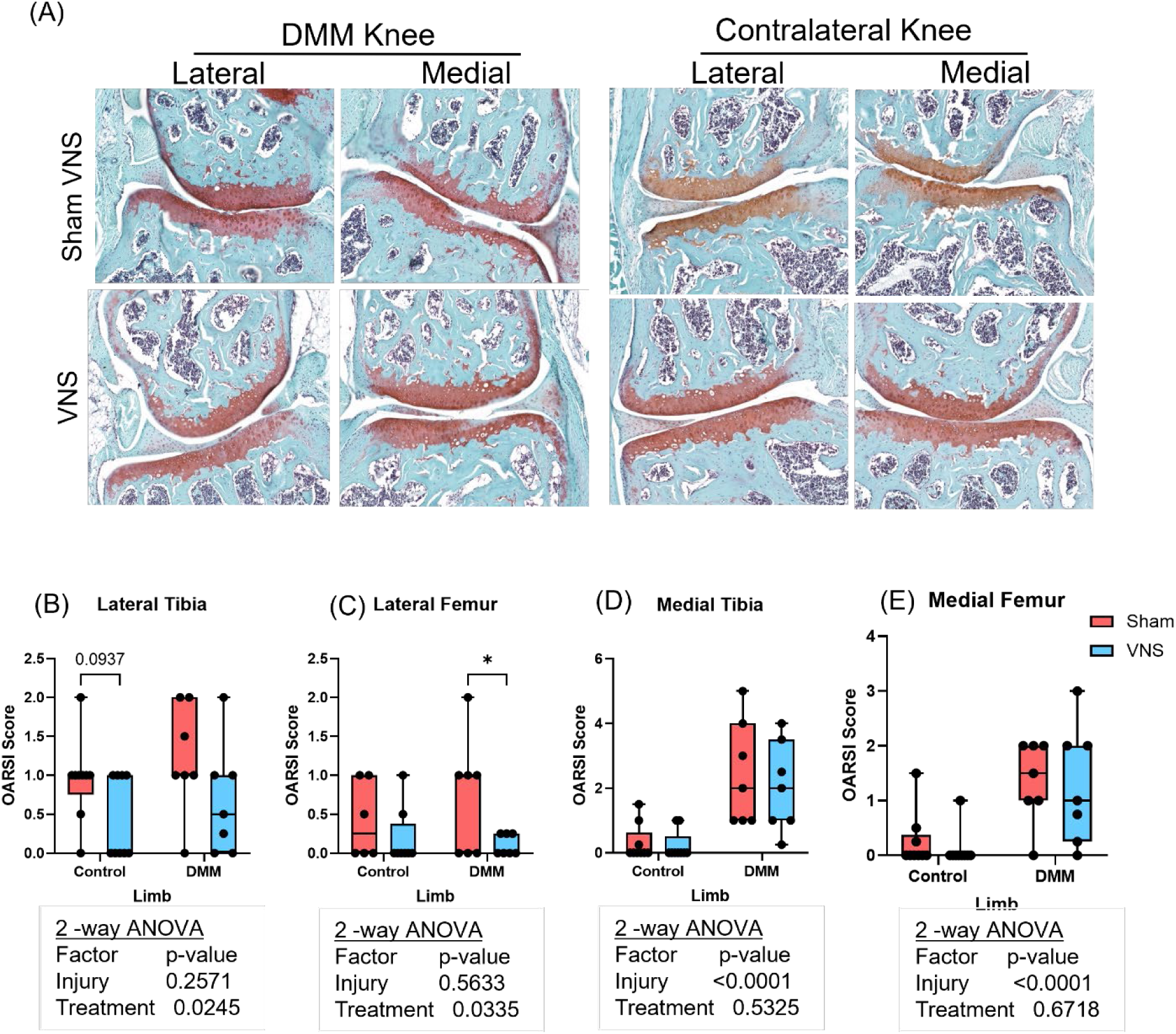
tVNS reduces knee damage in male mice with DMM-induced OA. (A) Representative safranin-O/fast green stained histology of the lateral and medial compartments from DMM and contralateral uninjured knees from tVNS and SHAM-treated mice. Below are graphs of OARSI scores for the SHAM and VNS groups of contralateral and DMM knees, specifically focusing on the (B) lateral tibia, (C) lateral femur, (D) medial tibia, and (E) medial femur. The lateral femur had a significantly reduced OARSI score after tVNS compared to SHAM treatment (C), with lateral tibia improvement approaching significance (B). Data are presented as mean ± SD, with n = 5 to 7; *p < 0.05. Two-way ANOVA followed by Fisher’s LSD multiple comparisons test was used to evaluate the effects of injury and treatment in the statistical analyses. Red is SHAM treated; Blue is tVNS treated.

### tVNS treatment leads to changes in serum inflammatory and pain biomarkers

Having established that tVNS treatment improves hyperalgesia and allodynia after acute ACL rupture and chronic weight-bearing pain in the DMM model, we then explored serum for biomarkers associated with these effects. We identified several serum cytokines associated with improved hyperalgesia and allodynia in female ACLR mice (Table 1 and Fig. 4A). Compared to sham treatment, tVNS induced significant reductions in IFN-γ, MIP-1β, IL-12p70 (Fig. 4A) and IL-3 levels (Table 1). No differences were seen in these cytokines in male mice. IL-1β was significantly reduced by tVNS in males and approached significance in females (Fig. 4A). tVNS treatment also significantly reduced Eotaxin and GM-CSF levels in males, while no differences in these cytokines were observed in females (Fig. 4A).

**Figure 4.**
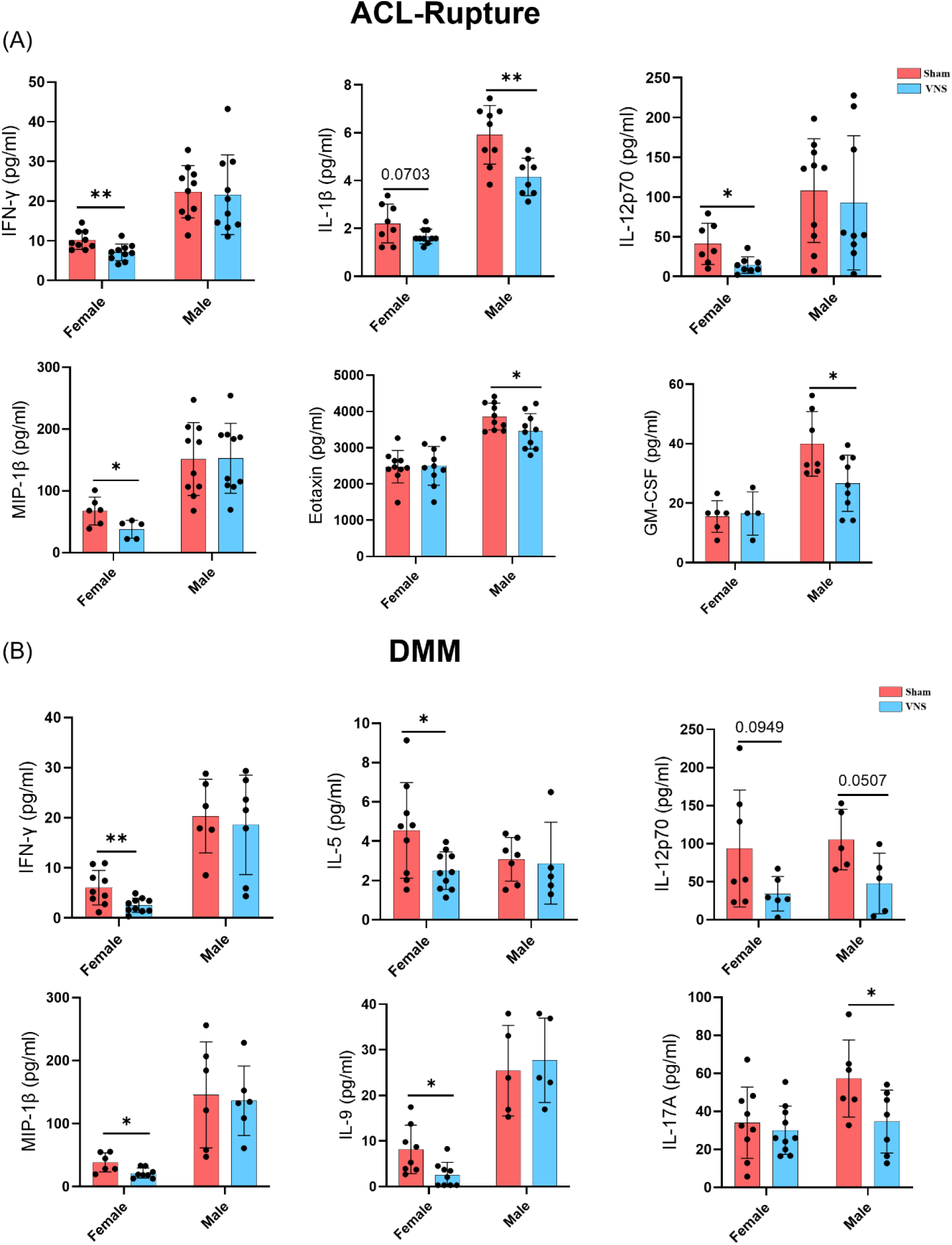
tVNS reduces serum pro-inflammatory cytokine and chemokine levels in PTOA. (A) Compared to SHAM stimulation, 2 weeks of tVNS treatment suppressed serum cytokines, including IL-1β, IFN-γ, MIP-1β, and IL-12p70 in female mice and 1L-1β, eotaxin, and GM-CSF levels in male mice. (B) Compared to SHAM treatment, 8 weeks of tVNS treatment in DMM mice decreased IFN-γ, MIP-1β, IL-5, and IL-9 levels in females and IL-17A in males. Data were presented as mean ± SD, with n = 5 to 10; *p < 0.05, **p < 0.01. Statistical analyses involved two-tailed unpaired t-tests or Mann-Whitney test comparing SHAM vs. tVNS in each model. Red is SHAM treated; Blue is tVNS treated.

**Table 1.**
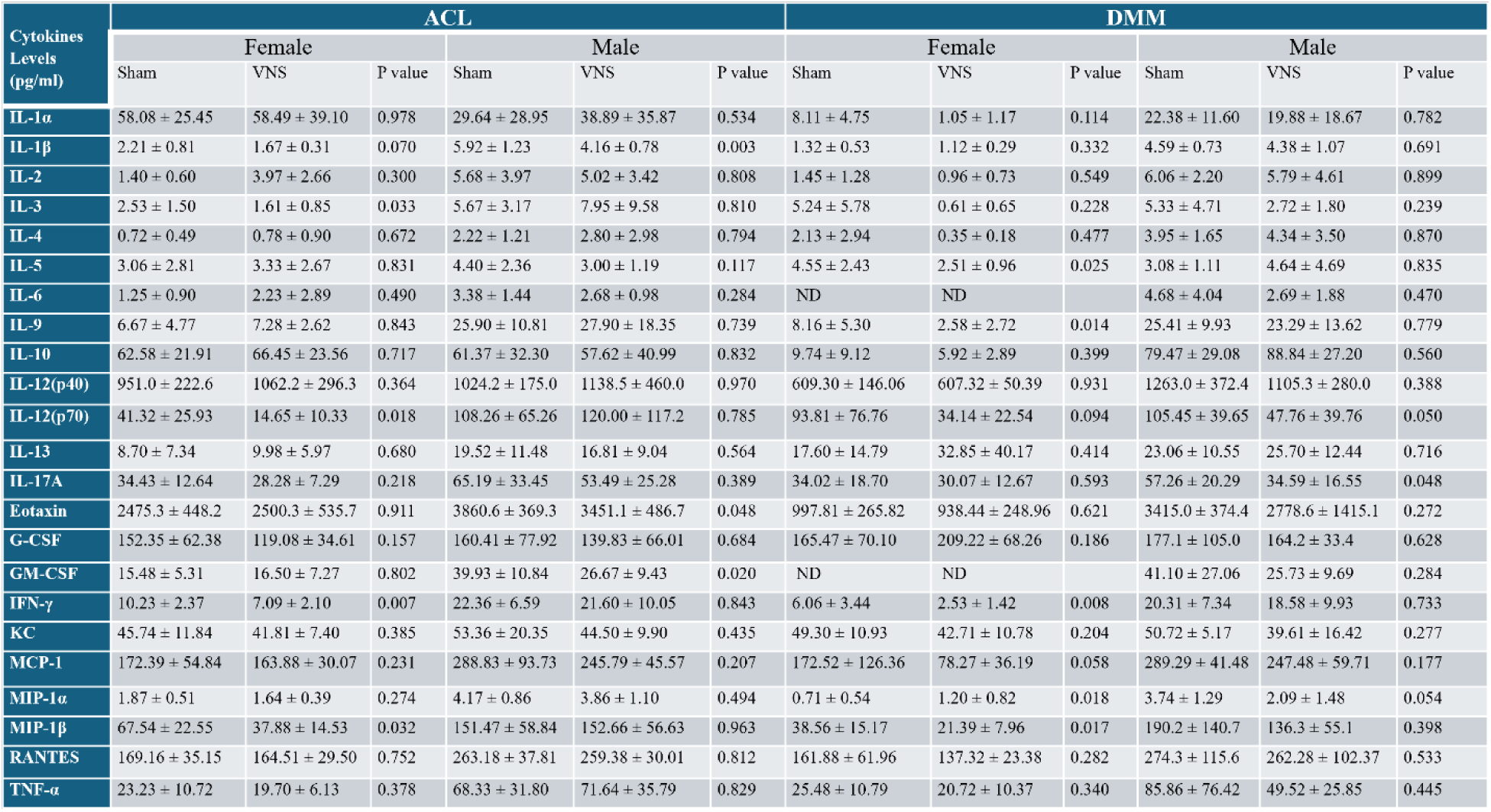
Serum cytokines modulated by tVNS in female and male ACL rupture and DMM-induced OA mice.

| Cytokines Levels (pg/ml) | ACL |  |  |  |  |  | DMM |  |  |  |  |  |
| --- | --- | --- | --- | --- | --- | --- | --- | --- | --- | --- | --- | --- |
|  | Female |  |  | Male |  |  | Female |  |  | Male |  |  |
|  | Sham | VNS | P value | Sham | VNS | P value | Sham | VNS | P value | Sham | VNS | P value |
| IL-1α | 58.08 ± 25.45 | 58.49 ± 39.10 | 0.978 | 29.64 ± 28.95 | 38.89 ± 35.87 | 0.534 | 8.11 ± 4.75 | 1.05 ± 1.17 | 0.114 | 22.38 ± 11.60 | 19.88 ± 18.67 | 0.782 |
| IL-1β | 2.21 ± 0.81 | 1.67 ± 0.31 | 0.070 | 5.92 ± 1.23 | 4.16 ± 0.78 | 0.003 | 1.32 ± 0.53 | 1.12 ± 0.29 | 0.332 | 4.59 ± 0.73 | 4.38 ± 1.07 | 0.691 |
| IL-2 | 1.40 ± 0.60 | 3.97 ± 2.66 | 0.300 | 5.68 ± 3.97 | 5.02 ± 3.42 | 0.808 | 1.45 ± 1.28 | 0.96 ± 0.73 | 0.549 | 6.06 ± 2.20 | 5.79 ± 4.61 | 0.899 |
| IL-3 | 2.53 ± 1.50 | 1.61 ± 0.85 | 0.033 | 5.67 ± 3.17 | 7.95 ± 9.58 | 0.810 | 5.24 ± 5.78 | 0.61 ± 0.65 | 0.228 | 5.33 ± 4.71 | 2.72 ± 1.80 | 0.239 |
| IL-4 | 0.72 ± 0.49 | 0.78 ± 0.90 | 0.672 | 2.22 ± 1.21 | 2.80 ± 2.98 | 0.794 | 2.13 ± 2.94 | 0.35 ± 0.18 | 0.477 | 3.95 ± 1.65 | 4.34 ± 3.50 | 0.870 |
| IL-5 | 3.06 ± 2.81 | 3.33 ± 2.67 | 0.831 | 4.40 ± 2.36 | 3.00 ± 1.19 | 0.117 | 4.55 ± 2.43 | 2.51 ± 0.96 | 0.025 | 3.08 ± 1.11 | 4.64 ± 4.69 | 0.835 |
| IL-6 | 1.25 ± 0.90 | 2.23 ± 2.89 | 0.490 | 3.38 ± 1.44 | 2.68 ± 0.98 | 0.284 | ND | ND | | 4.68 ± 4.04 | 2.69 ± 1.88 | 0.470 |
| IL-9 | 6.67 ± 4.77 | 7.28 ± 2.62 | 0.843 | 25.90 ± 10.81 | 27.90 ± 18.35 | 0.739 | 8.16 ± 5.30 | 2.58 ± 2.72 | 0.014 | 25.41 ± 9.93 | 23.29 ± 13.62 | 0.779 |
| IL-10 | 62.58 ± 21.91 | 66.45 ± 23.56 | 0.717 | 61.37 ± 32.30 | 57.62 ± 40.99 | 0.832 | 9.74 ± 9.12 | 5.92 ± 2.89 | 0.399 | 79.47 ± 29.08 | 88.84 ± 27.20 | 0.560 |
| IL-12(p40) | 951.0 ± 222.6 | 1062.2 ± 296.3 | 0.364 | 1024.2 ± 175.0 | 1138.5 ± 460.0 | 0.970 | 609.30 ± 146.06 | 607.32 ± 50.39 | 0.931 | 1263.0 ± 372.4 | 1105.3 ± 280.0 | 0.388 |
| IL-12(p70) | 41.32 ± 25.93 | 14.65 ± 10.33 | 0.018 | 108.26 ± 65.26 | 120.00 ± 117.2 | 0.785 | 93.81 ± 76.76 | 34.14 ± 22.54 | 0.094 | 105.45 ± 39.65 | 47.76 ± 39.76 | 0.050 |
| IL-13 | 8.70 ± 7.34 | 9.98 ± 5.97 | 0.680 | 19.52 ± 11.48 | 16.81 ± 9.04 | 0.564 | 17.60 ± 14.79 | 32.85 ± 40.17 | 0.414 | 23.06 ± 10.55 | 25.70 ± 12.44 | 0.716 |
| IL-17A | 34.43 ± 12.64 | 28.28 ± 7.29 | 0.218 | 65.19 ± 33.45 | 53.49 ± 25.28 | 0.389 | 34.02 ± 18.70 | 30.07 ± 12.67 | 0.593 | 57.26 ± 20.29 | 34.59 ± 16.55 | 0.048 |
| Eotaxin | 2475.3 ± 448.2 | 2500.3 ± 535.7 | 0.911 | 3860.6 ± 369.3 | 3451.1 ± 486.7 | 0.048 | 997.81 ± 265.82 | 938.44 ± 248.96 | 0.621 | 3415.0 ± 374.4 | 2778.6 ± 1415.1 | 0.272 |
| G-CSF | 152.35 ± 62.38 | 119.08 ± 34.61 | 0.157 | 160.41 ± 77.92 | 139.83 ± 66.01 | 0.684 | 165.47 ± 70.10 | 209.22 ± 68.26 | 0.186 | 177.1 ± 105.0 | 164.2 ± 33.4 | 0.628 |
| GM-CSF | 15.48 ± 5.31 | 16.50 ± 7.27 | 0.802 | 39.93 ± 10.84 | 26.67 ± 9.43 | 0.020 | ND | ND | | 41.10 ± 27.06 | 25.73 ± 9.69 | 0.284 |
| IFN-γ | 10.23 ± 2.37 | 7.09 ± 2.10 | 0.007 | 22.36 ± 6.59 | 21.60 ± 10.05 | 0.843 | 6.06 ± 3.44 | 2.53 ± 1.42 | 0.008 | 20.31 ± 7.34 | 18.58 ± 9.93 | 0.733 |
| KC | 45.74 ± 11.84 | 41.81 ± 7.40 | 0.385 | 53.36 ± 20.35 | 44.50 ± 9.90 | 0.435 | 49.30 ± 10.93 | 42.71 ± 10.78 | 0.204 | 50.72 ± 5.17 | 39.61 ± 16.42 | 0.277 |
| MCP-1 | 172.39 ± 54.84 | 163.88 ± 30.07 | 0.231 | 288.83 ± 93.73 | 245.79 ± 45.57 | 0.207 | 172.52 ± 126.36 | 78.27 ± 36.19 | 0.058 | 289.29 ± 41.48 | 247.48 ± 59.71 | 0.177 |
| MIP-1α | 1.87 ± 0.51 | 1.64 ± 0.39 | 0.274 | 4.17 ± 0.86 | 3.86 ± 1.10 | 0.494 | 0.71 ± 0.54 | 1.20 ± 0.82 | 0.018 | 3.74 ± 1.29 | 2.09 ± 1.48 | 0.054 |
| MIP-1β | 67.54 ± 22.55 | 37.88 ± 14.53 | 0.032 | 151.47 ± 58.84 | 152.66 ± 56.63 | 0.963 | 38.56 ± 15.17 | 21.39 ± 7.96 | 0.017 | 190.2 ± 140.7 | 136.3 ± 55.1 | 0.398 |
| RANTES | 169.16 ± 35.15 | 164.51 ± 29.50 | 0.752 | 263.18 ± 37.81 | 259.38 ± 30.01 | 0.812 | 161.88 ± 61.96 | 137.32 ± 23.38 | 0.282 | 274.3 ± 115.6 | 262.28 ± 102.37 | 0.533 |
| TNF-α | 23.23 ± 10.72 | 19.70 ± 6.13 | 0.378 | 68.33 ± 31.80 | 71.64 ± 35.79 | 0.829 | 25.48 ± 10.79 | 20.72 ± 10.37 | 0.340 | 85.86 ± 76.42 | 49.52 ± 25.85 | 0.445 |

Sham-stimulated DMM mice had significant increases in numerous cytokines (Table 1 and Fig.4B). Compared to sham treatment, 8 weeks of tVNS, a time point associated with improved weight-bearing, significantly reduced serum IFN-γ, IL-5, IL-9, and MIP-1β levels in females but not males (Fig. 4B). MIP-1α levels were also significantly reduced by tVNS in females, while in males, the reduction approached significance (Table 1). tVNS treatment significantly reduced IL-17A levels in male DMM mice (Fig. 4B). DMM female and male mice showed a decreasing trend in IL-12p70 with tVNS (Fig. 4B). Ther results show that tVNS treatment significantly reduces several inflammatory cytokines in both ACLR and DMM-induced PTOA.

### Association between allodynia and serum inflammatory biomarkers impacted by tVNS in ACL rupture

We performed a linear regression analysis of cytokine levels, von Frey withdrawal force, and percent weight bearing on ACL-injured limbs to determine the relationship between the identified serum biomarkers and pain outcomes. We found several significant findings. Specifically, the analysis yielded an R^2^ value of 0.28 (P=0.02) for IL-1β and 0.36 (P=0.006) for IFN-γ, demonstrating that variations in these cytokine levels explain 28% and 36% of the variability in von Frey thresholds, respectively (Fig. 5A). The inverse association observed indicates that as IL-1β and IFN-γ levels increase, von Frey threshold values decrease, signifying increased mechanical allodynia. Our data demonstrate that tVNS treatment decreases serum IL-1β and IFN-γ levels and increases von Frey thresholds in ACL-rupture female mice (Fig. 5A and 1B). Therefore, elevated IL-1β and IFN-γ levels are directly correlated with increased mechanical allodynia in ACL-injured limbs, and tVNS treatment reduces these, suggesting that IL-1β and IFN-γ could serve as biomarkers for mechanical allodynia and tVNS responses in ACL-injured female mice.

**Figure 5.**
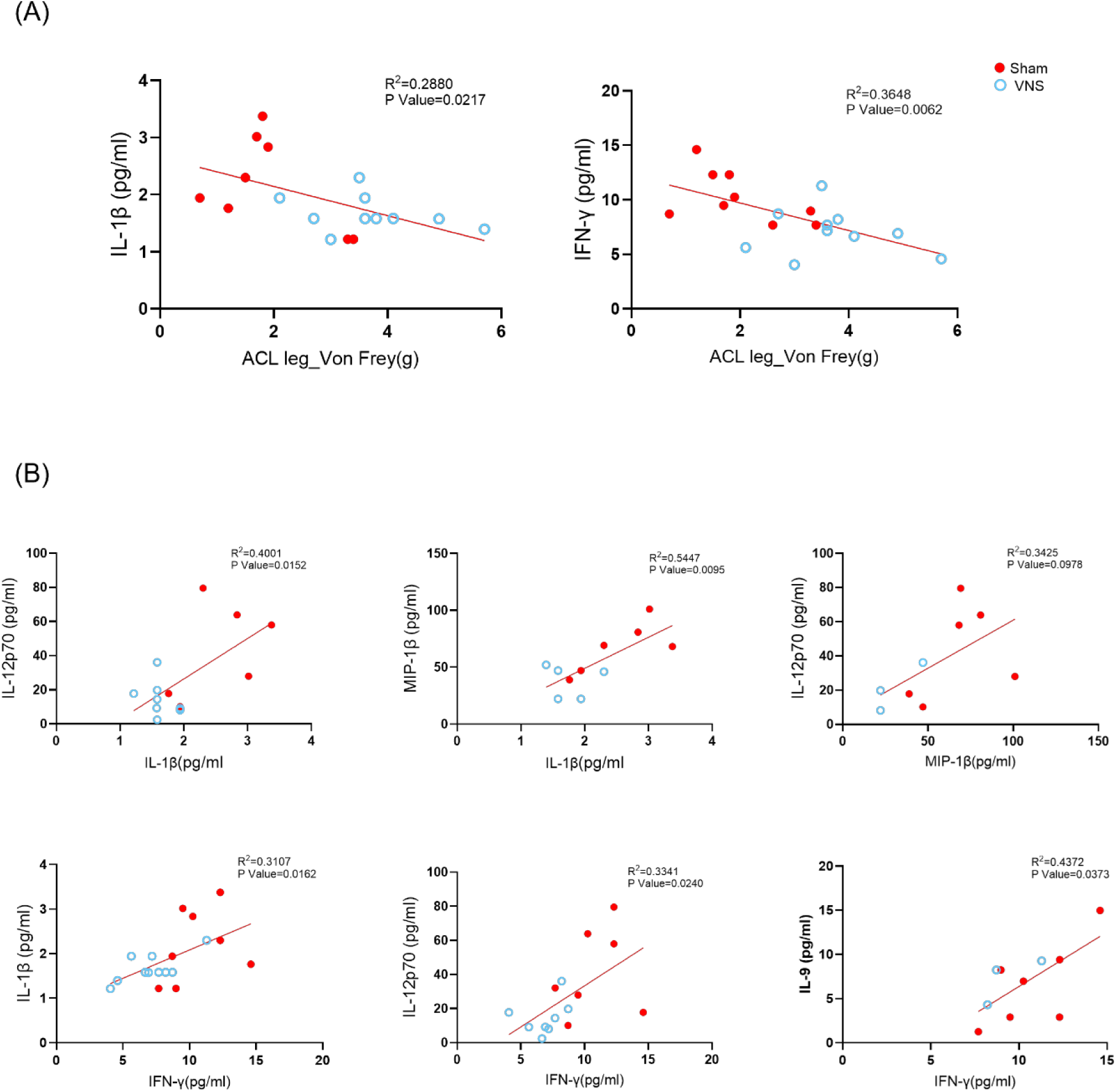
Linear associations between cytokines and von Frey responses improved with tVNS. (A) Associations between IL-1β or IFN-γ and von Frey withdrawal force. (B) Associations among tVNS modulated cytokines: IL-1β and IL-12p70, IL-1β and MIP-1β, MIP-1β and IL-12p70, IFN-γ and IL-1β, IFN-γ and IL-12p70, and IFN-γ and IL-9. The coefficient of determination (R²) and p values are presented in the respective graphs. Red circles are SHAM treated, and blue are tVNS treated.

Additional linear regression analysis revealed significant associations between the cytokine biomarkers that tVNS modulated. (Fig. 5B) IL-1β exhibited a moderate positive correlation with IL-12p70 (R^2^ = 0.40, P=0.02) and robust positive correlation with MIP-1β (R^2^=0.54, P=0.01), suggesting potential regulatory connections (Fig. 5B). Additionally, positive associations were found between IFN-γ and IL-1β (R^2^ = 0.31, P=0.02), IL-12p70 (R^2^ = 0.33, P=0.02), and IL-9 (R^2^ = 0.43, P=0.04) (Fig. 5B), hinting at cooperative mechanisms among these cytokines that tVNS impact. These findings emphasize the intricate interplay among these cytokines, suggesting potential regulatory mechanisms modulated by tVNS within the immune system.

### Associations between serum inflammatory biomarkers in DMM impacted by tVNS

The linear regression analysis conducted in DMM female mice revealed significant insights into the interplay of serum cytokines modulated by tVNS, showing notable associations. The robust correlation between IFN-γ and IL-12p70 is particularly striking, boasting an impressive coefficient of determination R² = 0.82 (P<0.001), suggesting a pivotal role in OA-related pain in female mice (S2). Equally noteworthy is the association between IFN-γ and IL-9, with an even higher R² value of 0.85 (P<0.001), underscoring their importance in modulating pain (S2). Furthermore, the correlation between IL-12p70 and IL-9 stands out with an R² value of 0.87, indicating a strong relationship between these cytokines (S2). IL-1β demonstrates correlations with IL-12p70, IL-5, and IL-9, with R² values of 0.45, 0.55, and 0.45, respectively, suggesting moderate to strong associations (S2). Similarly, IFN-γ exhibits notable associations with IL-1β, MIP-1β, and IL-5, with R² values of 0.51, 0.55, and 0.39, respectively (S2). MIP-1β demonstrates correlations with IL-5 and IL-9, supported by R² values of 0.64 and 0.43, respectively, while its association with IL-12p70 yields an R² of 0.46 (S2). Furthermore, IL-12p70 and IL-9 exhibit relatively weaker associations with IL-5, with R² values of 0.35 and 0.32, respectively (S2). These findings provide insights into the intricate cytokine network implicated in OA pathogenesis that are impacted by tVNS.

## Discussion

We have provided crucial preclinical studies showing the efficacy of noninvasive transcutaneous VNS (tVNS) in treating PTOA pain and reducing PTOA progression. We chose two PTOA models: the acute, inflammatory, and rapidly progressive ACL rupture model and the slowly progressive DMM model to assess the impact of tVNS on PTOA. We have shown that short-term tVNS improves mechanical hyperalgesia and allodynia in these two models, with more significant effects in female mice. The significance of these findings is highlighted by the fact that brief episodes (10 min) of stimulation exerted long-lasting anti-inflammatory effects and improved clinical outcomes in our rodent models, consistent with previous studies in animals and humans. (7, 14, 19). However, the minimum duration each day and number of tVNS sessions required to improve clinical outcomes remain to be optimized.

The sex differences we observed in PTOA pain responses are consistent with other studies showing enhanced pain responses in women and female animals despite less severe OA (20–23). Neuroimmune interactions, crucial for mechanical allodynia, differ between the sexes, with males predominantly showing microglial activation in the spinal cord, leading to increased neuronal excitability(24, 25), while females rely on adaptive immune cells, including T cells producing IFN-γ and IL-17A (26, 27). Furthermore, we have previously seen elevations in similar cytokines correlate with sex differences in OA histologic outcomes (28). Consistent with these studies, tVNS suppressed IL-1β in male mice, which is known to sensitize nociceptive neurons in the peripheral and central nervous systems, increasing nerve sensitivity and pain perception (29, 30). Elevations in IFN-γ and IL-12, type 1 cytokines known to correlate with increased pain and knee impairment in OA, were seen in female mice and reduced by tVNS. (31) In both models, tVNS reduced IFN-γ, MIP-1β, and IL-12p70 in females, suggesting a shared gender-specific pathway for suppressing pain and systemic inflammation. Thus, tVNS appears to reduce allodynia in female mice, possibly through adaptive immune cell mechanisms differing from male responses.

In the DMM model, tVNS significantly improved pain with weight bearing on the injured leg independent of sex. These findings were associated with reduced OARSI scores in the DMM model. However, tVNS did not affect weight bearing in the more severe ACLR PTOA model. tVNS did significantly reduce PTOA in both sexes (Fig. 2). This is a critical finding as other OA treatments have not shown to be DMOADs with the ability to reduce PTOA progression.

We speculate that decreases in OA progression with tVNS may be due to reductions in synovial inflammation and pro-inflammatory cytokines, IFN-γ, IL-1β, MIP-1β, IL-12, and IL-17, that promote cartilage degradation (32, 33). IL-1β promotes pro-inflammatory cytokines, activates matrix metalloproteinases (MMPs), and inhibits extracellular matrix (ECM) synthesis, leading to cartilage destruction (34). In chondrocytes, IFN-γ increases the transcription of key inflammatory mediators, including TNF-α, IL-6, and MMP-13 (35). IL-17 acts synergistically with IL-1β and TNF-α, enhancing inflammatory responses and promoting MMPs and aggrecanases that degrade the cartilage matrix (34). Thus, these cytokines play crucial roles in cartilage degradation, emphasizing their importance in OA progression.

Our study has several limitations. The changes in serum cytokines seen in our models suggest tVNS regulates specific populations of immune cells. IFNγ and IL-12 are type 1 cytokines produced by CD4+ T cells and are known to be enriched in OA synovium (36, 37). T helper 9 cells are upregulated in OA and produce IL-9, which drives mast cell expansion and contributes to cartilage breakdown (38–40). We speculate that tVNS modulates these cells in the peripheral blood or locally within the joint, a subject for future studies. We have previously shown that sex-linked differences in C57BL6/J OA susceptibility are partly mediated by the gut microbiome, leading to systemic immune cell changes (28). We have not yet investigated whether the gut microbiome contributes to tVNS responses in our OA models, but other studies show that tVNS can alter the gut microbiome in irritable bowel syndrome (41). Additional studies are required to determine whether tVNS modulates dorsal root ganglia or the central nervous system, including pain-processing areas like the brainstem, which may further influence pain perception and relieve chronic pain conditions in a sex-dependent manner. These additional investigations will also provide insight into the sex-specific tVNS responses seen in our OA models, forming the basis for future studies in humans with acute ACL ruptures and in OA. Based on our findings, women may be more responsive to tVNS than men in these settings.

In summary, we present the first evidence that tVNS is a promising therapeutic approach to reduce PTOA pain and progression in two preclinical models. These results strongly support pilot studies of tVNS in acute ACL-tears and chronic OA.

## Contributors

MBH, MJ, SS, and TG conceived the experiment.SY, DD, LM, TC, JL, SS, EP, VI, GD, and MB planned and conducted the experiments. SY, VI, SS, JL, TG, MBH, and MJ performed data analysis. SY, MBH, MJ, and TG contributed to interpreting results. SY and MBH wrote the manuscript. All authors provided critical feedback and helped shape the research, analysis, and manuscript. All authors approved the final manuscript.

## Acknowledgements

Funding source for this work was supported by the I01BX006046 VA BLR&D Merit Review Award (MBH, MJ, TG, SS) and the Presbyterian Health Foundation Award (MBH, MJ, TG, SS).

---

**Sup Figure 1.**
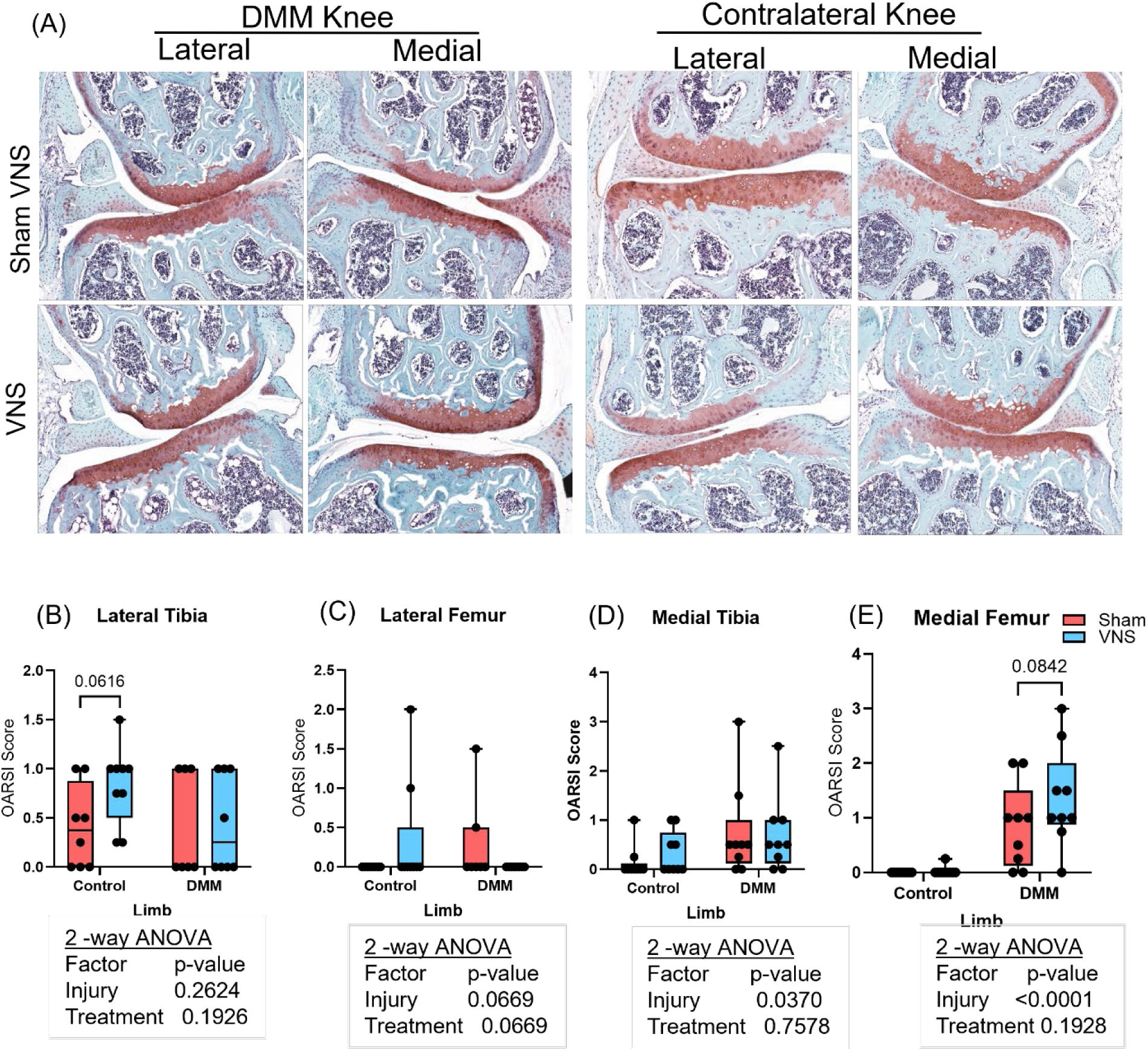
DMM in female mice induces minimal OA that tVNS does not modulate. (A) Representative safranin-O/fast green stained histology of the lateral and medial compartments from DMM and contralateral uninjured knees from tVNS and SHAM-treated female mice. Below are graphs of OARSI scores for the SHAM and tVNS groups of contralateral and DMM knees, specifically focusing on (B) lateral tibia, (C)lateral femur, (D) medial tibia, and (E) medial femur. Data are presented as mean ± SD, with n = 5 to 9. Two-way ANOVA followed by Fisher’s LSD multiple comparisons test was used to evaluate the effects of injury and treatment in the statistical analyses. Red is SHAM treated; Blue is tVNS treated.

**Sup Figure 2.**
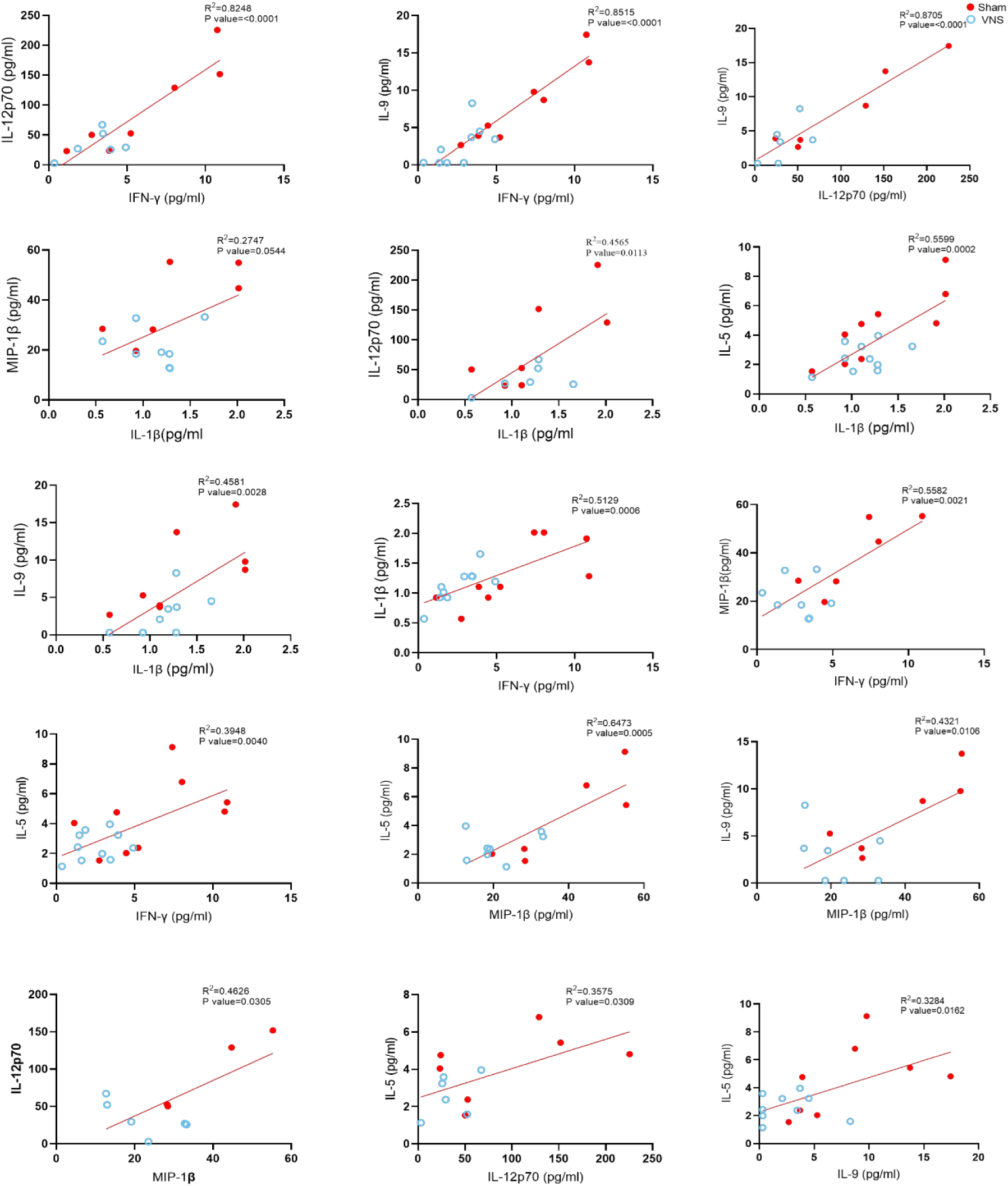
Significant linear associations between tVNS modulated cytokines in DMM Female Mice. The strongest linear associations are observed between IFN-γ and IL-12p70 (R² = 0.82). IFN-γ and IL-9 (R² = 0.85), and IL-12p70 and IL-9 (R² = 0.87). Additionally, there are significant associations, including IL-1β with IL-12p70, MIP-1β, IL-5, and IL-9. The coefficient of determination (R²) and p-value are presented in the respective graphs. Red circles are SHAM treated, and blue are tVNS treated.

